# Evolution is exponentially more powerful with frequency-dependent selection

**DOI:** 10.1101/2020.05.03.075069

**Authors:** Artem Kaznatcheev

## Abstract

Valiant [1] proposed to treat Darwinian evolution as a special kind of computational learning from statistical queries. The statistical queries represent a genotype’s fitness over a distribution of challenges. And this distribution of challenges along with the best response to them specify a given abiotic environment or static fitness landscape. Valiant’s model distinguished families of environments that are “adaptable-to” from those that are not. But this model of evolution omits the vital ecological interactions between different evolving agents – it neglects the rich biotic environment that is central to the struggle for existence.

In this article, I extend algorithmic Darwinism to include the ecological dynamics of frequency-dependent selection as a population-dependent bias to the distribution of challenges that specify an environment. Thus, extended algorithmic Darwinism suggests extended statistical queries rather than just statistical queries as the appropriate model for eco-evo dynamics. This extended algorithmic Darwinism replaces simple invasion of wild-type by a mutant-type of higher scalar fitness with an evolutionary game between wild-type and mutant-type based on their frequency-dependent fitness function. To analyze this model, I develop a game landscape view of evolution, as a generalization of the classic fitness landscape approach.

I show that this model of eco-evo dynamics on game landscapes can provide an exponential speed-up over the purely evolutionary dynamics of the strict algorithmic Darwinism. In particular, I prove that the Parity environment – which is known to be not adaptable-to under strict algorithmic Darwinism – is adaptable-to by eco-evo dynamics. Thus, the ecology of frequency-dependent selection does not just increase the tempo of evolution, but fundamentally transforms its mode. This happens even if frequency-dependence is restricted to short-time scales – such short bursts of frequency-dependent selection can have a transformative effect on the ability of populations to adapt to their environments in the long-term.

Unlike typical learning algorithms, the eco-evo dynamic for adapting to the Parity environment does not rely on Gaussian elimination. Instead, the dynamics proceed by simple isotropic mutations and selection in finite populations of just two types (the resident wild-type and invading mutant). The resultant process has two stages: (1) a quick stage of point-mutations that moves the population to one of exponentially many local fitness peaks; followed by (2) a slower stage where each ‘step’ follows a double-mutation by a point-mutation. This second stage allows the population to hop between local fitness peaks to reach the unique global fitness peak in polynomial time. The evolutionary game dynamics of finite populations are essential for finding a short adaptive path to the global fitness peak during the second stage of the adaptation process. This highlights the rich interface between computational learning theory, analysis of algorithms, evolutionary games, and long-term evolution.

---

**E**volution is an algorithm. As such, evolution is subject to the same constraints of computational complexity as all algorithms [1–3]. But evolution is not *any* algorithm. It is a kind of local search algorithm acting on the signal of fitness. Depending on if we treat fitness as an intrinsic property of organisms that is independent of the distribution of other organisms (frequency-independent fitness) or as a function of that distribution (frequency-dependent fitness) changes the power of the algorithm of evolution. By better understanding the kinds of algorithms that are evolutionary, we can better understand the power and limits of evolution in nature.

In this article, I will show that adding ecological interactions – conceived of as frequency-dependent selection – to Valiant [1]’s frequency-independent evolution has a qualitatively transformative effect on evolutionary dynamics. Ecology provides a fundamentally different computational resource for evolution that allows exponential speed-ups in the time it takes to adapt to certain environments. This is not a speed-up in the rate of evolution, but a fundamental shift in how evolution computes and thus which ‘problems’ it can ‘solve’: i.e. which kinds of environments populations can or cannot become well-adapted to. Ecology does not make evolution take steps faster along an exponentially long adaptive path but instead ecology helps evolution find a short path. Thus, ecology does not just increase the tempo of evolution, but fundamentally transforms its mode. To demonstrate this, I will focus on a family of highly non-linear environments – the Parity problem defined more formally in Section 3 – that evolution on its own (i.e. without ecology) provably cannot adapt to.

Valiant [1]’s strict algorithmic Darwinism (described in Section 1) formalizes many of our intuitions about evolution and allows for the analysis of a richer set of evolutionary dynamics than most traditional models in biology. But it lacks an important feedback mechanisms found in nature: eco-evolutionary feedback. I will introduce this feedback as part of the extended algorithmic Darwinism in Section 2.

From first glance, it might seem like ecology would not have a drastic effect on the adaptive power of evolution: especially in settings when ecological dynamics are much faster than evolutionary dynamics. This is probably why this feedback was often ignored or minimized in early models of the modern evolutionary synthesis. Consider, for example, the *E. Coli* long-term evolution experiment that has been tracking genetic changes in 12 initially identical populations of asexual bacteria since 24 February 1988 – recently passing the time-scale of 73,500 generations. Here it is known that if we want to look at short-term coexistence of organisms then frequency-dependent fitness matters [4, 5]. However, the strength of the frequency-dependent selection is on the order of 1-5%. Thus, when biologists look at the long-term dynamics – the repeated replacement of types by new types – then they find it adequate to focus on just the frequency-independent component of fitness [6–8]. In other words, *in practice*, when evolutionary biologists want to predict long-term dynamics, they find it reasonable to ignore the relatively weak frequency-dependent selection [9, 10]. In this article I ask: is ignoring weak frequency-dependent selection also reasonable *in theory* ?

Even in the modern eco-evo literature, much of the work is motivated by settings where ecology and evolution are on similar timescales [11, 12]. If ecological interactions are on a much faster timescale than evolution, it seems like ecological dynamics can be “absorbed” into a single update of evolutionary dynamics. This might make one “step” of evolution faster or slower, but running faster in no particular direction or along an exponentially long path is not helpful. Thus, it feels intuitive that short-bursts of ecological interaction within a single step of evolution cannot fundamentally change what sort of environments populations can adapt to. This intuition is wrong.

Short bursts of weak frequency-dependent selection can transform long-term dynamics of evolution.

Even if ecological dynamics happen on a much faster timescale than evolution, I will show that the feedback between ecology and evolution due to brief bursts of frequency-dependent selection can help eco-evolutionary dynamics find short adaptive paths that purely evolutionary dynamics cannot find. I will show this by concentrating on if populations can find the peak of the needle-in-a-haystack fitness landscape corresponding to the Parity environment. In Section 3, I will define the Parity environment and rehearse a standard hardness argument to show that under strict algorithmic Darwinism, the population cannot adapt to this environment in polynomial time (regardless of which evolutionary dynamic it follows). But in Section 5, I will show that the simple mutation-limited strong-selection dynamics from Section 4 are sufficient for the population to adapt to the parity environment in polynomial time under extended algorithmic Darwinism.

From the perspective of the design and analysis of algorithms, this paper’s main contribution is a careful analysis of a particular algorithm on a particular family of inputs. In Section 5 I prove the following:

## Theorem 1.

*The Parity family of environments is adaptable-to in the framework of extended algorithmic Darwinism*

In particular, the Parity family of environments is adaptable-to by simple mutation-limited strong-selection dynamics in finite populations with just isotropic point-mutations and double mutations. This is a sharp contrast to the typical learning algorithms for Parity that tend to rely on Gaussian elimination.

Like the dynamics themselves, my analysis of these evolutionary dynamics proceeds in two parts. In Proposition 3, I show that evolution quickly converges to a set 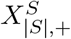 of genotypes that are local peaks with respect to point-mutations (Proposition 3) through a fast phase of point mutations. In Proposition 4, I show that this is followed by a slow phase of a biased-random walk by double-mutations among the set 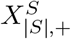 to the global fitness peak. The overall time is *O*(*n/µ*) fixations (where *µ* is the mutation rate) or *O*(*n*^7^) death-birth events (Proposition 5).

From the perspective of computational learning theory, Theorem 1 seperates the complexity class corresponding to strict algorithmic Darwinism from the complexity class corresponding to extended algorithmic Darwinism:

## Corollary 2.

*Extended algorithmic Darwinism with frequency-dependent selection is more computationally powerful than strict algorithmic Darwinism*.

This exponential speed up from ecology is due to the ability of the ecological dynamics of the evolving population to bias the challenge/example distribution in such a way that creates a new second-order gradient for evolution to follow. This allows for a much richer set of concept classes to be learnable by evolution (or in more biological terminology, a richer set of environments that are adaptable-to). For upper bounds, this means replacing the tight CSQ-learnable upper bound on strict algorithmic Darwinism [13] by the potentially loose extended-SQ-learnable (or PAC + membership queries) [14] upper bound on the extended algorithmic Darwinism. I leave it as an open problem to see exactly how much more computationally powerful eco-evolutionary dynamics can be in comparison to just evolutionary dynamics.

## 1 Strict algorithmic Darwinism: abiotic/prebiotic interactions

To represent what evolution can achieve just by natural selection on frequency-independent fitness, I will focus on Valiant’s model of strict algorithmic Darwinism [1, 15]. In my presentation, I will change some of the terminology to more closely correspond to that of the biologists. In particular, ‘evolvability’ has a rich meaning in the biology literature that does not always correspond to Valiant [1]’s use of the word. So my primary change will be to use statements like ‘a family of environments is adaptable-to’ instead of ‘a concept class is evolvable’ and correspondingly I will refer to the model as ‘(strict) algorithmic Darwinism’ instead of ‘evolvability’. These are only changes in terminology for talking about the same mathematical model.

As a running biological motivation throughout the article, I will ground the model in the primordial evolution of early life. This interpretation is not central to the mathematics and if you have a different preferred biological system for the contrast of frequency-independent vs frequency-dependent fitness then feel free to substitute it in your reading.

### Abiotic environment ((*D, f*))

Let us consider an abiotic world filled with a primordial soup of complex polymers. For simplicity, let us represent each possible polymer by some string *z* ∈ *Z* = {0, 1} ^*n*^ where *n* is our primary size parameter for asymptotic arguments. Let 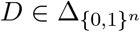 be the particular probability distribution of these polymers in the soup (from the set of all possible distributions on the possible polymers; 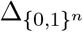). We can think of each polymer *z* as a ‘challenge’ sampled from *D*. Let us suppose that for each polymer *z* there are exactly two ways {0, 1} to interact with it and one of these ways is ‘correct’ and the other is ‘incorrect’ – where the correct way liberates useful energy and the incorrect way spoils the polymer. Let the ideal function *f* : {0, 1}^*n*^ → {0, 1} hold the ‘correct’ response to each polymer – we can think of this *f* as the physical laws that govern the polymers in our hypothetical world. Thus, the pair (*D, f*) can be seen as a representation of a particular abiotic environment.

### Genotypes (*x*) and phenotypes (*g*_*x*_)

A primordial organism has to encounter these abiotic challenges and respond. We can summarize the organism’s responses as a (behavioral) phenotype *g*_*x*_ : {0, 1}^*n*^ → {0, 1} that is encoded by a genotype *x* ∈ *X* = {0, 1}^*m*^. In general, there will be many particular token organisms that all have the same genotype *x* and phenotype *g*_*x*_.

### Fitness (*w*_*x*_ and *ŵ*_*x*_)

We can imagine a specific token organism of type *x* that is floating in this primordial soup and bumping into polymers as sampling a challenge *z* ∈ *D*. If the response of the organism to this challenge (as given by *g*_*x*_(*z*)) is the ‘correct’ response (as given by *f* (*z*)) then the organism gets, as payoff, a small reproductive reward (if *g*_*x*_(*z*) = *f* (*z*)) and otherwise it gets no reward (if *g*_*x*_(*z*) ≠ *f* (*z*)).

Of course, evolution itself does not act on tokens but rather on types [16, 17]. Evolution can only respond to statistical properties of many individuals (tokens) of the same type, each encountering many individual challenges sampled from *D*. Moreover, evolution cannot access arbitrary statistical properties of *D* – as Kearns [18]’s statistical query model allows – but instead has to use the particular statistical signal of (type) fitness. Formally, we can define 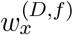 as the true fitness of *x* in environment (*D, f*):

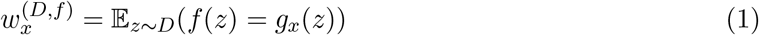

which I will also write as *w*_*x*_ when the environment (*D, f*) is obvious from context. A real population, however, will not have direct access to *w*_*x*_ and instead will have to rely on taking the average fitness over a random sample *L* ∼ *D* of *lM* many challenges, where *M* is the size of the population and *l* is the number of challenges per token. I will use 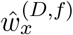 to represent the empirical fitness estimate of the true fitness 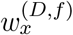. Throughout the analysis, it will be useful to upper bound the error between the empirical and true fitness value by a tolerance *α* as 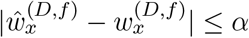.

The exact value of *α* depends on the population size *M* and the number of challenges per token *l*. If we wanted to ensure a tolerance of *α* with high probability then we should set 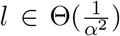. The above is defined for a physics *f* that is deterministic and error-free, but with a reasonable number of extra random queries, this tolerance can be ensured even if *f* is sometimes corrupted by noise (i.e. an organism having a small probability of getting a fitness bonus even if it responded to the challenge incorrectly or not getting fitness even if it responded correctly) [18]. Thus, as is typical with statistical queries, we can use the same analysis of adaptability for both standard environments and noisy environments.

### Fitness landscape (*w* or *W*) view of strict algorithmic Darwinism

It is also possible to express strict algorithmic Darwinism in the long established metaphor of fitness landscapes [19] by setting *w*(*x*) = *w*_*x*_. A point *x* ∈ *X* is said to be a local peak if all its point-mutation (single bit flip) neighbours (*N* (*x*)) are non-improving (i.e. ∀*y* ∈ *N* (*x*) *w*(*x*) ≥ *w*(*y*)) and a global peak if all points are non-improving. For multiplicative fitness measures that are exponential functions of payoffs – like number of offspring – we can instead use 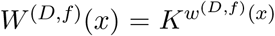 for some constant *K* > 1 or equivalently *W* ^(*D,f*)^(*x*) = exp(*βw*^(*D,f*)^(*x*)) for some strength of selection *β* [20].

### *Adaptable-to* families of environments ((𝒟, ℱ))

Since algorithmic biology is after asymptotic results, we are usually concerned not with a particular abiotic environment (*D, f*) but a family of environments (𝒟, ℱ) defined over a class of possible distributions 𝒟 and set of potential ideal functions ℱ = {*f*_*n,s*_} (i.e. concept class) indexed by some natural number *n*, index *s*. Borrowing an idea from his earlier model of probably-approximately correct learning [21], Valiant [1] defines an environment (𝒟, ℱ) as *adaptable-to* if given an arbitrarily small probability 0 ≤ *δ <* 1*/*2 and arbitrarily small approximation rate 0 ≤ *ϵ <* 1*/*2 there exists an evolutionary dynamic that in time polynomial in *n*, |*s*|, ln 1*/δ*, and 1*/ϵ* with probability 1 − *δ* gets the population to a genotype *x* such that Pr_*z*∼*D*_(*g*_*x*_(*z*) = *f* (*z*)) ≥ 1 −*ϵ* for any *D* ∈ 𝒟 and any *f*_*n,s*_ ∈ ℱ.

In the language of fitness landscapes, a family of environments (𝒟, ℱ) is adaptable-to if evolutionary dynamics can find the global fitness peak or get to a type that has fitness *E* close to it in the corresponding family of fitness landscapes. Or more formally, if for every (*D*_*n*_, *f*_*n,s*_) with probability 1 − *δ* the population reaches a genotype *y* with 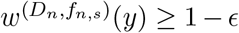 in time polynomial in *n*, |*s*|, ln 1*/δ*, and 1*/ϵ*. Note that this is different from Kaznatcheev [2] and Kaznatcheev, Cohen, and Jeavons [3]’s focus on local peaks.

Time to find local or global peaks is the most convenient way to connect computational complexity to evolution but it is not the only measure of computational power or efficiency that we could use to analyze evolution as an algorithm. For example, a biologist might be interested in the mean rate of increase in fitness or rate of adaptation [6–8, 22, 23], even without expecting an optimum to be reached. However, in the idealized case of the parity fitness landscapes that I focus on later in this article, there will be only a few discrete fitness levels. So, in the case of the parity fitness landscape, not finding an optimum will mean remaining in the same fitness plateau of *w* = 1*/*2. Thus, for this article, I will define families of environments as adaptable-to or *easy* if those environments’ global fitness peak’s fitness can be approximated within *E* in polynomial time and not adaptable-to or *hard* otherwise. To understand the computational power of algorithmic Darwinism, I will be interested in understanding the complexity class that contains all the families of environments that are adaptable-to.

The computational power of strict algorithmic Darwinism is well understood. Feldman [13] showed that strict algorithmic Darwinism is computationally equivalent to a subset of Kearns [18]’s statistical query model that is restricted to making only correlational queries (CSQ). In other words, a family of environments (𝒟, ℱ) is adaptable-to if and only if the concept class F is CSQ-learnable over the distributions 𝒟 = {*D*_*n*_}. This means that the parity function over the uniform distribution – discussed in Section 3 – is not adaptable-to. But since the parity function *is* PAC-learnable, Valiant [1] used this as an argument for why learning is more powerful than evolution.

## 2 Extended algorithmic Darwinism: biotic/ecological interactions

In this article, I extend the strict Algorithmic Darwinism of Valiant [1] to accommodate frequency-dependent fitness. The goal is not to build the most realistic model but to make the smallest deformation that introduces frequency-dependent fitness. Let me sketch this in terms of the prebiotic soup of early life.

### Abiotic distribution (*D*_*A*_)

For Valiant, the distribution *D* of environmental challenges is not affected by the resident population: all hypothetical populations in the same world would experience the same distribution of challenges. Consider, for example, challenges as macromolecules in the prebiotic soup: we can imagine the action of the sun on the early chemistry of the Earth producing a consistent supply of random polymers for our organisms to encounter. This would be a sort of constant abiotically generated distribution *D*_*A*_ of challenges.

### Full distribution with biotic component (*D*(*ρ*))

But the abiotic distribution is not the only source of polymers in the environment: other organisms can also be a source of polymers. Thus, we can imagine the distribution *D* as partitioned into two parts, an abiotic part *D*_*A*_ that is independent of other organisms in the environment and a biotic part *D*_*B*_(*ρ*) that is a function of the evolving population *ρ* (here I will take *ρ* ∈ Δ_*X*_ as the distribution of types in the population). This gives us *D*(*ρ*) = (1 − *b*)*D*_*A*_ + *bD*_*B*_(*ρ*), where *b* ∈ [0, 1] measures the strength of the biotic component. In this extended algorithmic Darwinism, Valiant’s strict model would correspond to *b* = 0 and the fitness signal would be a kind of statistical query. By incorporating frequency-dependent selection with *b* > 0, we move from SQ to extended statistical queries. I focus the rest of the discussion on a family of landscapes where *b* is a small but polynomial fraction (i.e. *b* ≈ 1*/n*^*O*(1)^). This fraction *b* cannot be exponentially small since that would correspond to a “non-biased” distribution and “non-biased” distributions are known to be insufficient for learning Parity in the extended statistical query model [14].

In a general application, the space of possible challenge strings *Z* and the space of genotypes *X* might not be the same. In that case, *D*_*B*_: Δ_*X*_ → Δ_*Z*_ needs to map from distributions on the space of possible genotypes (Δ_*X*_) to distributions on the space of possible challenge strings (Δ_*Z*_). We can view this mapping as specifying the ecological part of the evolutionary algorithm, since it would encode what kind of challenges certain organisms tend to present to other organisms. But in this article, I will focus on when *X* = *Z* = {0, 1}^*n*^ and set *D*_*B*_ to be identity thus making *D*(*ρ*) = (1 − *b*)*D*_*A*_ + *bρ*. In the case of the protocells in a prebiotic soup, we can imagine *D*(*ρ*) implemented as all protocells trying to constantly copy their own genotypes but due to their primordial nature and poor lipid membranes, constantly shedding some of these copies without always successfully duplicating. When one cell floats past another, it then encounters this cloud of shed copies as new challenges. Thus, this would bias the abiotic distribution *D*_*A*_ towards the distribution of protocells *ρ* by an amount *b* that will depend on various aspects of the environmental and population structure.

### Game landscapes (*ω*)

For the sake of this article, I will focus on matrix games as a model of frequency-dependent fitness. For strict algorithmic Darwinism, we could have imagined the *w* of a fitness landscape as a very long vector in ℝ^*X*^ with entries indexed by *x* ∈ *X* and specified by Equation 1. Similarly, for the extended algorithmic Darwinism, I will build the (matrix) game landscapes by considering a very large matrix *G* ∈ ℝ^*X*×*X*^ with entries given by:

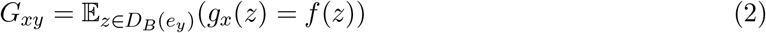

where *e*_*y*_ ∈ Δ_*X*_ is the distribution with all weight at type *y* (i.e. the unit vector in direction *y*). When *X* = *Z* and *D*_*B*_ is the identity, we can specify fitness functions for the game landscape as:

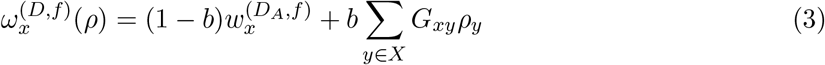

### Darwinian engine that powers evolution

I visualize this whole model as the Darwinian engine that powers evolution in Figure 1. The Darwinian engine is made of two cycles that together change the distribution of genotypes. On the top, we have the genesis of new variants via the mutation cycle. On the bottom, we have the struggle for existence via the development-ecology-selection cycle. The game landscape acts as a summary of the contribution of development (the mapping from genotype to phenotype) and ecology (from the distribution of phenotypes to fitness) that I described in this section. This is the heart of algorithmic Darwinism and is what specifies the environment that the population is adapting to – the problem instance, for a computer scientist. The other two arrows (selection and mutation) specify the algorithm of evolution. If we want to prove intractability results then we want to reason about arbitrary (polynomial time computable) selection and mutation functions. But since I aim for a surprising tractability result, I will describe in Section 4 the specific evolutionary algorithm of strong-selection weak-mutation dynamics [24, 25].

**Figure 1:**
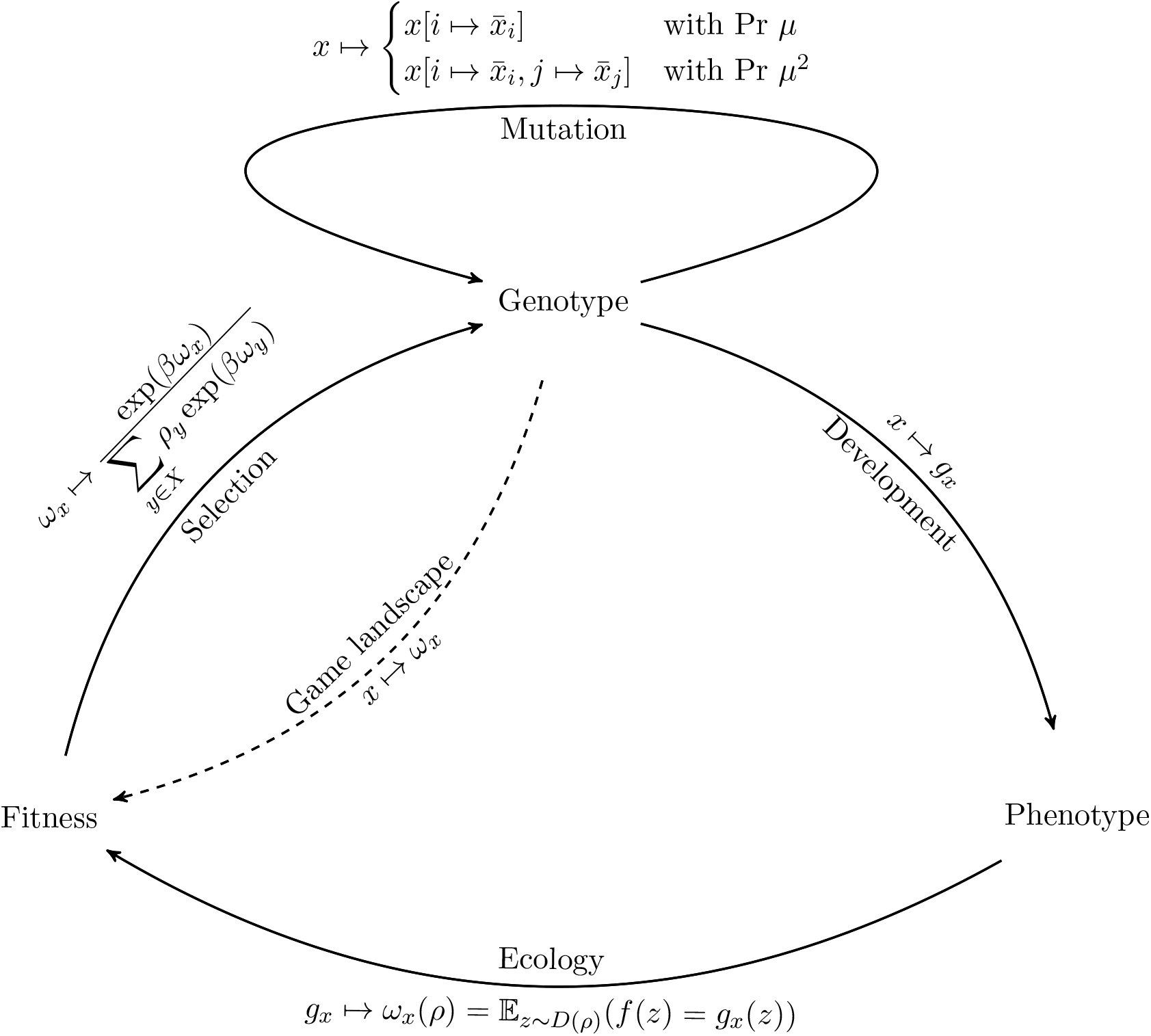
Darwinian engine powering eco-evolutionary dynamics. The top cycle captures the genesis of new variants and the bottom cycle captures the struggle for existence. Each edge is labeled by both its conceptual role (in text) and its specific realization in the ecologically extended algorithmic Darwinism (in equations). The two edges (selection and mutation) pointing into the genotype node specify the algorithm of evolution that adapts the population. In general, we could consider arbitrary polynomial time algorithms for these edges, but for the tractabiltiy of the Parity environments, I focus on the specific algorithm of strong-selection weak-mutation dynamics in a Moran process with single and double mutations as presented in Section 4. The other edges (development, ecology, and the out-direction of mutation) specify the environment (or ‘problem instance’) that the population is adapting-to. I give a general overview of this in Section 2 and discuss the specific family of Parity environments in Section 3.

## 3 Problem and representation: parity environments

We can now look at the specific ‘problem’ or family of environments that I consider in this article and define some terminology to help with our analysis. The canonical example of a hard environment for strict algorithmic Darwinism is the Parity environment [1].

### Parity environments (*f*_*n,s*_)

For each size *n*, the challenges come from the uniform distribution *U*_*n*_: i.e., every string in {0, 1}^*n*^ is as likely as every other string and has an equal probability of 1*/*2 of being encountered. Each environment of size *n* is indexed by a hidden set *S* of salient bits or – equivalently – an indicator string *s* ∈ {0, 1}^*n*^ that has *s*_*i*_ = 1 if *i* ∈ *S* and 0 otherwise. The correct response to a challenge *z* ∈ {0, 1}^*n*^ is to return the parity of the bits *z*_*i*_ for *i* ∈ *S* (i.e. 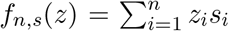 mod 2). In other words, to succeed organisms have to pay very careful attention to all the positions in *S* and ignore all other positions.

We might be interested handling noisy functions where an organism has a small probability of getting a fitness bonus even if it responded to the challenge incorrectly or not getting fitness even if it responded correctly. We can always set our tolerance parameter *α* low enough to account for this kind of misclassification noise in *f*. Thus, I can treat *f*_*n,s*_ as not noisy but, without loss of generality, the main results will still hold for noisy-Parity.

### Parity is hard for strict algorithmic Darwinism

After defining SQ-learning, Kearns [18] showed that Parity over the uniform distribution is not efficiently learnable with statistical queries, no matter how clever we are with our algorithm or representation of the hypothesis class. Since any family of environments that is adaptable-to by strict algorithmic Darwinism has to be an SQ-learnable concept class, Valiant [1] gave Parity environments as the prototypical hard family of environments: a family of environments that are not adaptable to by any evolutionary dynamics compatible with strict algorithmic Darwinism. Since the parity function *is* PAC-learnable, Valiant [1] uses this as an argument for why learning is more powerful than evolution. Although it is important to note that noisy-Parity is conjectured to not even be PAC-learnable [26, 27].

### Representing phenotypes as genotypes

The above hardness result for strict algorithmic Darwinism is true regardless of how the phenotypes *g* are encoded as genotypes. Giving a specific mapping between genotypes and phenotypes (*X* → (*Z* → {0, 1})) and the distribution of genotypes and distribution of examples (Δ_*X*_ → Δ_*Z*_) can be seen as specifying the representation part of an evolutionary algorithm in the extended algorithmic Darwinism framework.

For the sake of an intuition about this hardness result, and as preparation for the easiness result in Section 5, I want to consider a specific simple representation of phenotypes as genotypes. Let the genotype space be the same as the example space *X* = *Z* = {0, 1}^*n*^ with the identity mapping between them. This will let us see the genotype *x* ∈ {0, 1}^*n*^ as the indicator set for the parts of the environment that the organism pays attention to, thus defining the phenotype *g*_*x*_ which on encountering challenge *z* produces 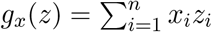 mod 2.

### Head and tail of genotypes

During the analysis in Section 5, I will call the part of the genotype that overlaps with the hidden set of salient bits (*x*[*S*]) the *head* of *x* (and use *λ*_*s*_(*x*) = |*x*[*S*]| for the size of the head), and the other part the *tail* (and use *τ*_*s*_(*x*) = |*x*[[*n*] − *S*]| for the size of the tail). See Figure 2a, for a specific visual example of how head and tail are defined given the salient string *s* and a genotype *x*.

**Figure 2:**
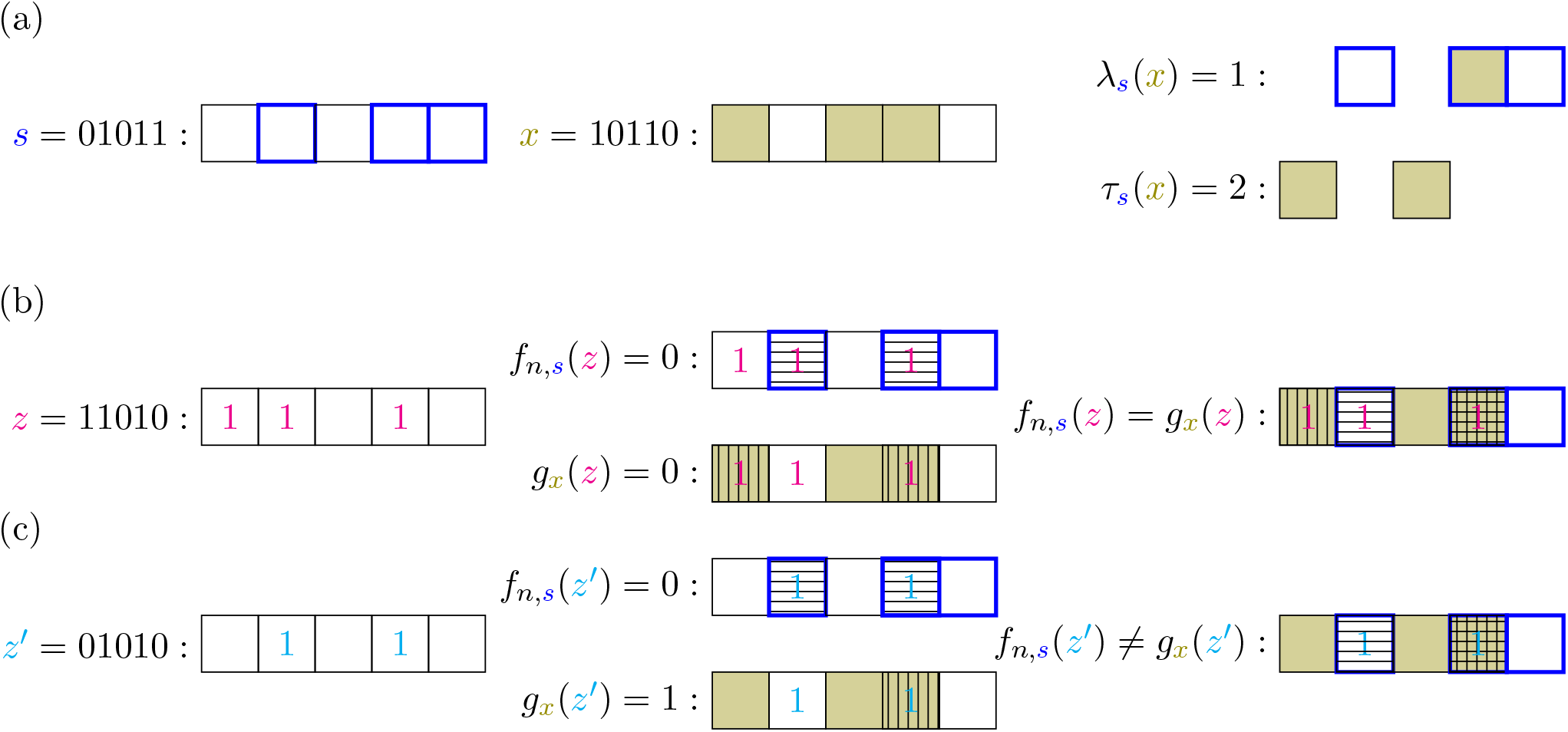
Example organism with genotype *x* in parity environment with salient bits *s* (a) responding to two different challenges *z* (b) and *z*^′^ (c). Consider an example parity environment on *n* = 5 bits (shown as line of 5 boxes) with salient set *S* = {2, 4, 5} (thus, salient bits *s* = 01011), shown as blue-bordered boxes. An organism with genotype *x* = 10110 (shown as olive-filled boxes) then has a head of size *λ*_*s*_(*x*) = 1 (due to bit 4) and tail of size *τ*_*s*_(*x*) = 2 (due to bits 1, 3) in this environment. We can look at two sample challenge strings *z* = 11010 (b) and *z*^′^= 01010 (c) in this environment. The correct response to the challenges is given by *f*_*n,s*_ and calculated based on the number of salient bits in the challenge, i.e. the number of 1 bits of the challenge that are in the salient set (highlighted as boxes with horizontal bars). If the number of salient bits in the challenge is even (i.e.,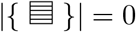 mod 2) then the correct response is 0, if it is odd (i.e.,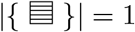 mod 2) then the correct response is 1. The response of the organism with genotype *x* is given by its phenotype *g*_*x*_ and calculated based on the number of bits in the challenge that the organism considers as salient, i.e. the number of 1 bits in the challenge that are also 1s in *x* (highlighted as boxed with vertical bars). If the number of bits in the challenge that the organism considers as salient is even (i.e., 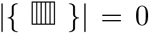 mod 2) then the organism responds with 0, if it is odd (i.e.,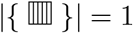 mod 2) then the organism responds with 1. The resulting fitness effect on the organism depends on if the response was correct (for example in (b), where *f*_*n,s*_(*z*) = *g*_*x*_(*z*)) or incorrect (for example in (c), where *f*_*n,s*_(*z*^′^) ≠ *g*_*x*_(*z*^′^)). If the response was correct then the organism gets a fitness benefit and if the response was incorrect then the organism gets no benefit. It is important to note, that sometimes the organism can be correct by accident. In (b), for example, the correct response to *z* is 0 and the organism responds with 0, but the reasons are different. The correct response is 0 because {2, 4} ⊆ *S* (and |{2, 4}| = 0 mod 2) but the organism responds as 0 because it thinks that {1, 4} is salient. Since only the response – and not the reason – matters, the organism with genotype *x* would get a fitness benefit from challenge *z*.

For every 0 ≤ *k* ≤ |*S*|, let 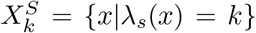 be the set of genotypes with a head of size *k*. I will say that a genotype *x* has a full head if *λ*_*s*_(*x*) = |*S*| (i.e., if 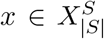). A genotype *x* has an *even-tail* if it has an even number of 1s in the tail (i.e. if *τ*_*s*_(*x*) = 2*k* for some integer *k*) and an *odd-tail* otherwise. Let us call the set of all even-tailed genotypes as 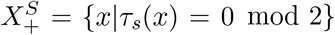 and the set of all odd-tailed genotypes as 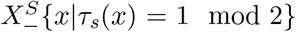. Combining this with the notation from above, we can let 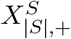 be the set of all full-head even-tail genotypes. This set will be important in our analysis of evolutionary dynamics in Section 5. It will also be useful to use the shorthand *λτ* (*x*) = (*λ*_*s*_(*x*), *τ*_*s*_(*x*)) for the head-tail coordinates of a genotype *x*.

See Figures 2(b,c), for a specific visual example of how a particular *g*_*x*_ acts on two particular samples and the kind of reward this yields in a particular environment with salient bits *s*.

### Parity fitness landscape (*w*^*s*^)

This representation of phenotypes by genotypes specifies a fitness landscape (*w*^*s*^) corresponding to the environment (*U*_*n*_, *f*_*n,s*_) that shows the intuition for why the Parity environment is hard: fitness simply does not provide much information about *s*. For a genotype *x* = *s* we have *w*^*s*^(*s*) = 1, but for any other genotype *x≠s*, we have that *w*^*s*^(*x*) = 1*/*2. This is because given any string *x*, there are exactly 2^*n*−1^ strings that share an even number of bits in common with *x* and also 2^*n*−1^ strings that share an odd numbers of bits in common with *x*. Thus, exactly half the time *g*_*x*_(*z*) evaluated to 0 and the other half the time to 1 – i.e. *g*_*x*_(*z*) agrees with *f*_*n,s*_(*z*) exactly half of the time.

For convenience, we can write down *w*^*s*^ as a polynomial or simple VCSP-instance [3]:

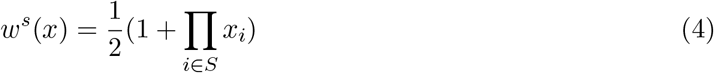

From this, we can easily see that the fitness landscape is a needle-in-the-haystack landscape: one giant lowlands at fitness 1*/*2 with a single peak hiding at genotype *s* out of the 2^*n*^ possible genotypes. In this case, where the fitness landscape is not providing any information, mutation limited strong selection dynamics of the kind outlined in Section 4 will with high probability take an exponential number of steps.

### Biological plausibility as the problem of selective attention

It is important to note that the biological plausibility of the Parity environment is not relevant for the separation of strict algorithmic Darwinism from the extended algorithm Darwinism. If we want to demonstrate that frequency-dependent fitness makes evolution qualitatively more powerful as an algorithm then finding any family of environments that is hard for strict algorithmic Darwinism and easy for the extended algorithmic Darwinism will do. Parity is just a particularly convenient choice for analysis because it is so well studied in the computational learning theory literature.

Equation 4 also highlights why parity functions are a theoretically important class of functions for biologists. Since the parity functions form a nice basis for all functions from {0, 1}^*n*^ → ℝ, any fitness function can be decomposed as a sum of parity functions. And, for example, looking at the largest |*s*| among the parity functions in this decomposition will tell us the highest-order epistasis in the landscape. So although we might not consider a needle-in-a-haystack fitness landscape with a single high-order epistasic component as very plausible, they can be seen as the building blocks for more ‘realistic’ fitness landscapes. Thus, having a deep theoretical understanding of when individual parity functions can and cannot be adapted-to can be useful to understanding how evolution proceed on other kinds of fitness functions.

With that said, although Parity might not seem like a very biologically plausible environment, I think it actually idealizes an important problem that many organisms are likely to face. The Parity environment, divides the sensory/informational world of the organisms into two parts: an important part *S* and an irrelevant part [*n*]−*S*. The correct response to an environmental challenge is extreme sensitive to the important part (with any one bit flipping the output) and completely insensitive to the irrelevant part (with the output unaffected by bit flips). But the irrelevant part still gives the organisms opportunities to attend to it (thinking that it matters to what should be done when it does not) and by so doing potentially make mistakes. The difficulty of the Parity environment then is not in the computation of the exclusive-OR of the various bits (which is not common in biological systems, but certainly possible for a single cell [28, 29]), but in separating the set of important stimuli from the set of irrelevant stimuli. Thus, in the context of simple biological systems with minimal cognition [30], the Parity environment can be seen as an extreme version of the problem of selective attention.

## 4 Algorithm: Mutation-limited dynamics for game landscapes

In general, the strict and extended algorithmic Darwinism allow for a rich set of possible evolutionary dynamics. And when we prove hardness results in these frameworks, we establish that no evolutionary dynamic compatible with the framework is capable of adapting-to some environment. However, when we are establishing a tractability result, it is useful to consider a particular evolutionary dynamic that is simple and biologically plausible.

### Monomorphic and briefly polymorphic populations

In the case of traditional fitness landscapes, for the sake of analysis, biologists often consider a mutation-limited population [25]. This population is nearly always monomorphic (i.e. made up of a number of tokens that are all of a single type) except when a new mutant is invading, at which point the population is briefly polymorphic between the resident and invader type. The invading mutant is usually a point-mutant of the resident type for timescales on the order of 1*/µ* where *µ <<* 1 is the small mutation rate. If the resistant cannot be invaded by any point-mutant then it is possible for a double (or more distant) mutant to arise, although the timescales for these events scale as (1*/µ*)^2^ where *µ <<* 1 is the small mutation rate. After a successful invasion, the population is again monomorphic. This corresponds to the extreme limit when ecological feedback can act only briefly during a single step of evolutionary dynamics.

In the case of a traditional fitness landscape, strong selection mutation-limited dynamics only allows a genotype *y* to invade and replace a genotype *x* if *w*(*y*) ≥ *w*(*x*). In strict algorithmic Darwinism, a mutant invades if it is strictly fitter (*w*(*y*) − *w*(*x*) > *t*) or nearly-neutral [31] (|*w*(*y*) − *w*(*x*)| ≤ *t*). But the story is more interesting for a game landscape and – instead of a simple threshold – it becomes useful to explicitly study a finite population of size *M*.

### Fitness of resident (*ω*_*x*_) and mutant type (*ω*_*y*_) as a function of number of mutants (*M*_*y*_)

Since we are focused on the limit where at most two genotypes *x, y* ∈ *X* co-exist at a time, we can track the state of the population with a single integer *M*_*y*_ ∈ [*M*] for the number of invading individuals of type *y*. For simplicity, I will assume a well mixed population so that every protocell has the same chance of interaction with every other protocell. This means that in a polymorphic population, a cell of type *x* has a probability of *M*_*y*_*/*(*M* − 1) of interaction with a cell of type *y* and a probability of (*M* − *M*_*y*_ − 1)*/*(*M* − 1) of interaction with a cell of type *x*. Similarly, a cell of type *y* has a probability of (*M*_*y*_ − 1)*/*(*M* − 1) of interaction with a cell of type *y* and a probability of (*M* − *M*_*y*_)*/*(*M* − 1) of interaction with a cell of type *x*. This transforms Equation 3 as a function of *M*_*y*_ into:

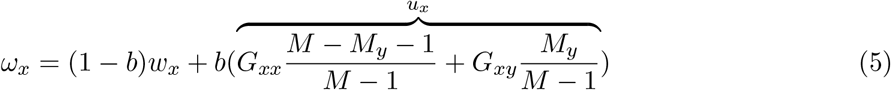

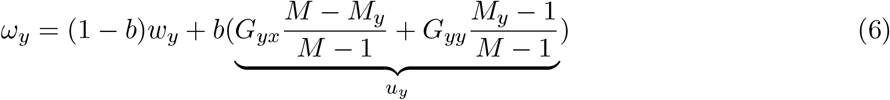

When analyzing evolutionary games, it is useful to look at the (whole) gain function (*γ*_*yx*_ = *ω*_*y*_ −*ω*_*x*_) of switching from *x* to *y* [32, 33] – also known as the invasion fitness:

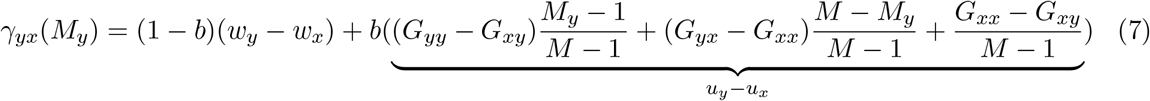

and when *w*_*y*_ = *w*_*x*_, it is useful to focus on just the component corresponding to the game gain function (*u*_*y*_ − *u*_*x*_). Since *f* has a boolean range, we have that *G*_*xx*_, *G*_*xy*_, *G*_*yx*_, *G*_*yy*_ ∈ {0, 1} and thus only 16 possible game gain functions, as shown in Figure 3. These kind of two strategy matrix games are well studied in evolutionary game theory and can even be measured directly in microscopic experimental systems [17, 34].

**Figure 3:**
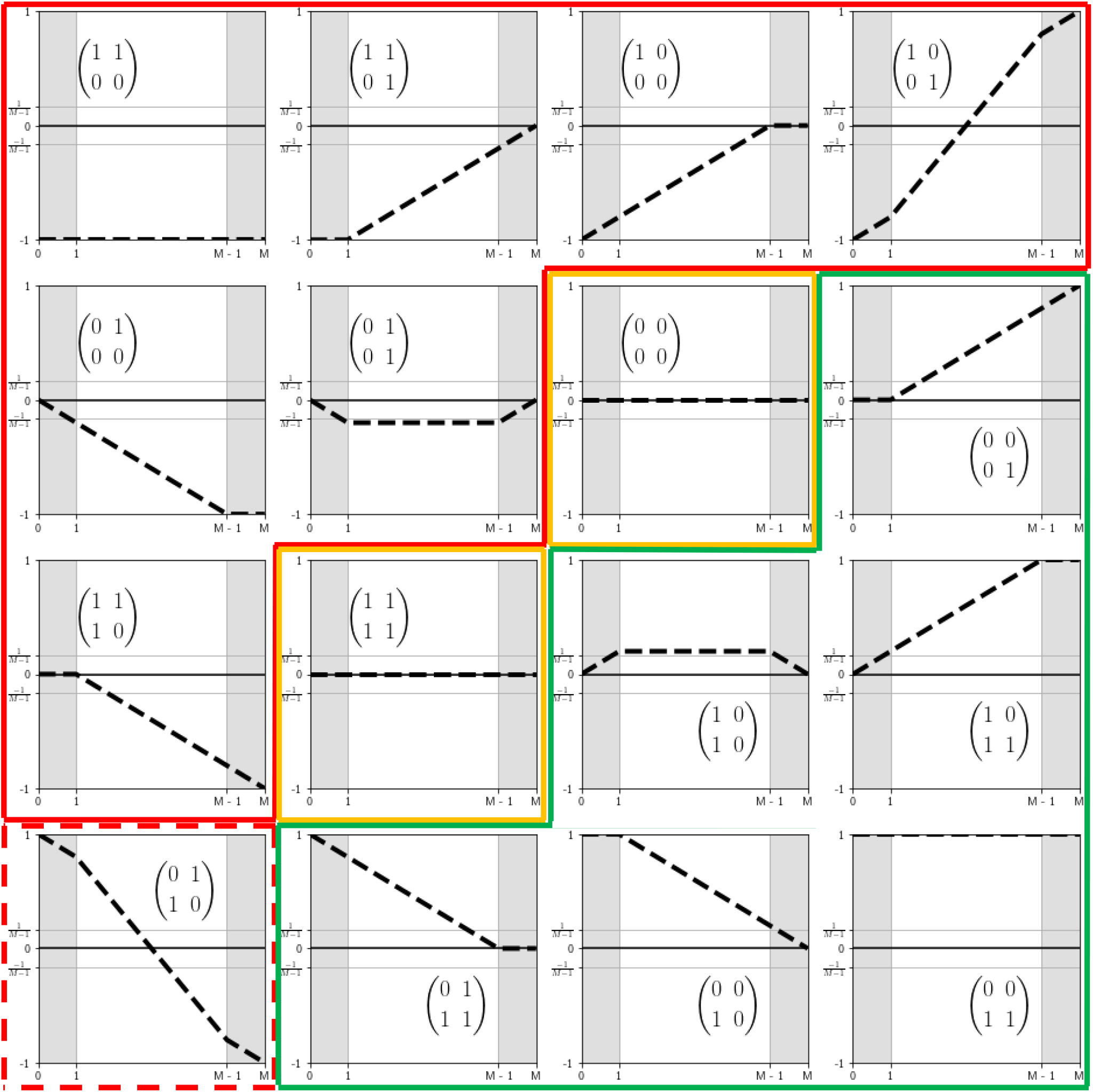
Possible game gain functions for game landscapes of a Boolean environment: For each graph, the x-axis is the number (*M*_*y*_) of invaders of type *y* and the y-axis is the game gain function (*u*_*y*_ −*u*_*x*_). The game of each graph is inset as the matrix 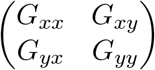. The possible invasion dynamics under strong selection (*β* > *M*) are given by the colour-coding: green means *y* can invade *x* with probability greater than some constant *ϕ*^*^; yellow means *y* can invade *x* through random drift with probability 1*/M* ; red means *y* cannot invade *x* (solid red: nearly 0 probability of fixation; dotted red: high probability, but exponentially long time to fixation).

### Fixation probability of mutant against resident (*ϕ*_*yx*_)

Let us use the Moran process [20, 35] based on fitnesses *ω*_*x*_ and *ω*_*y*_ to update the number of invaders *M*_*y*_. The Moran process is a standard model of invasion of a wild-type by a mutant-type in evolutionary biology, and proceeds as follows: (1) Randomly select an individual *m* proportional to it’s multiplicative fitness *e*^*βω*^ [20]; (2) Uniformly select an individual *c* to be replaced; (3) Replace *c* by *m* and repeat the process until the population is monomorphic. This defines a Markov chain on the state space [*M*] with transition probabilities given by:

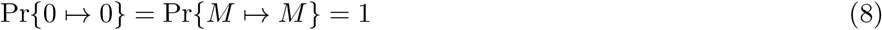

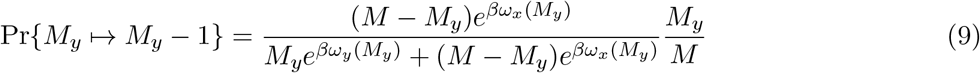

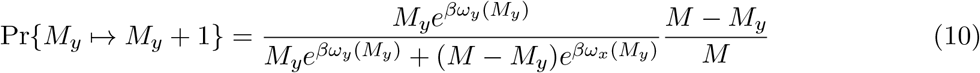

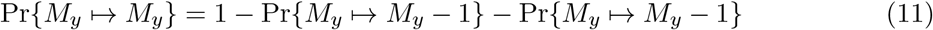

From this, we can calculate the fixation probability *ϕ*_*yx*_ of *y* (i.e. the probability that the above Markov chain starting at *M*_*y*_ = 1 ends at the absorbing state of *M*_*y*_ = *M*) as:

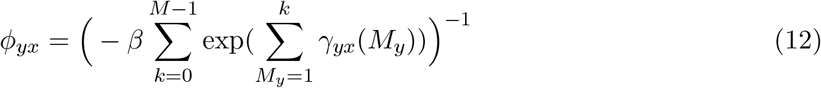

### Strong selection

I will focus on strong selection (*β* > *M*) and when the game contributes a detectable amount of the fitness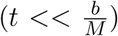. In that case, it is easy to classify the possible game regimes of the extended algorithmic Darwinism in Figure 3 into 4 classes:

1. In the green region, invasion is very likely: *ϕ*_*yx*_ > *ϕ*^*^ for a constant *ϕ*^*^ independent of the rest of the parameters;
2. In the yellow region, invasion is possible but not likely (the neutral and nearly-neutral mutation regime): *ϕ*_*yx*_ = 1*/M* ;
3. In the red region, invasion is effectively impossible: *ϕ*_*yx*_ *< e*^−*M*^ ;
4. In the dotted red region, invasion is likely but the fixation time is exponential in *M* (this regime will not occur in the particular case of the parity game landscape that I analyze in this article).

With a fitness landscape, it can be helpful to replace the numeric structure by just the structure of allowed adaptive moves to get the fitness graph [2, 3, 36, 37]. Similarly, with a game landscape, we can imagine it as a set of nodes {0, 1}^*n*^ connected by the four types of edges outlined above. If type (4) edges are not encountered, my separation of time-scales between ecology and evolution – much like similar separations in the adaptive dynamics literature [12, 38, 39] – allows us to continue to view a population as occupying a single vertex. But unlike the simple continuous landscapes of adaptive dynamics, game landscapes allow us to also preserve the rich combinatorial structure and discrete mutations that in fitness landscapes allows for computationally hard environments [2].

## 5 Parity is easy for extended algorithmic Darwinism

As we look at the mutation-limited dynamics in the Parity environment in the ecological setting, it is important to look at not only the static fitness function of *w*^*s*^(*x*) (or 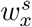) but also at the two-strategy matrix game 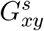 (which I will abbreviate as *G*_*xy*_ when *s* is clear from context) that becomes:

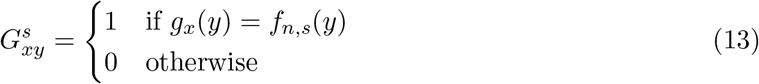

Note that when *x* = *s*, 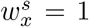 and for all *y* ∈ {0, 1}^*n*^,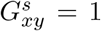. Thus, just like the fitness landscape, the game landscape also has a single unique global fitness peak at *s*. To see how evolution on the Parity game landscape finds the *s* peak, we have to focus on how a mutation-limited strong selection dynamics navigates the great low fitness plateau where for all *x, y* ∈ {0, 1}^*n*^ − {*s*}, 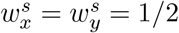. Here, we can focus on just the biotic contribution and the game gain functions from Figure 3 that account for every possible invader. This lets us analyze the evolutionary dynamics of adapting to Parity. I will show that these dynamics proceed by two stages, a fast stage where each step is a point-mutation and thus happens on the time-scale of 1*/µ* (Proposition 3) and a slow stage where each step is a double-mutation and thus happens on the time-scale of 1*/µ*^2^ (Proposition 4).

### Proposition 3.

*Given any parity environment f*_*n,s*_, *genotypes with a full-head and even tail (i*.*e*., *genotypes in* 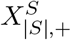) *are evolutionary stable under point-mutations and, starting from any genotype, SSWM dynamics in the extended Algorithmic Darwinism framework converge to a genotype in* 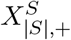*after O*(*n*) *many fixations of point-mutations*.

*Proof*. Even in the monomorphic case, there is a deformation as we shift from a fitness to a game landscape. In particular, we see the great low-fitness plateau at fitness 1*/*2 split into two parts:

- Consider any even-tail genotype *x*: in this case, we have *G*_*xx*_ = 1. Thus, in a monomorphic population of all-*x*, we have the biotic contribution of *u*_*x*_ = 1 and the same abiotic contribution of *w*_*x*_ = 1*/*2 giving a total fitness of *ω*_*x*_ = (1 + *b*)*/*2.
- Consider any odd-tail genotype *y*: in this case, we *G*_*yy*_ = 0). Thus, in a monomorphic population of all-*y*, we have the biotic contribution of *u*_*y*_ = 0 and the same abiotic contribution of *w*_*y*_ = 1*/*2 giving a total fitness of *ω*_*y*_ = (1 − *b*)*/*2.

Since any odd-tail genotype is adjacent to [*n*] − *S* many even-tail genotypes, the set of odd-tailed genotypes is a valley and not a plateau since any *y* with an odd-tail can be invaded by any adjacent *x* with the same head *y*[*S*] = *x*[*S*] but an even-tail. In particular, the games between *y* (with *λτ* (*y*) = (*λ*, 2*k* − 1)) and *x* depend on if *x*’s tail is small and thus a subset of *y*’s (i.e., *λτ* (*x*) = (*λ*, 2*k* − 2)) or if if it is larger and thus a superset (i.e., *λτ* (*x*) = (*λ*, 2*k*)):

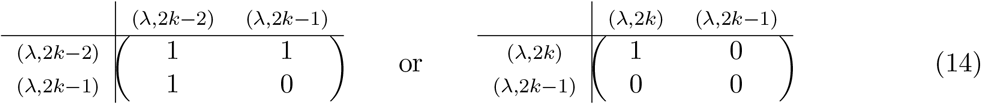

The resulting slice of the game landscape can be seen visually in Figure 4a.

**Figure 4:**
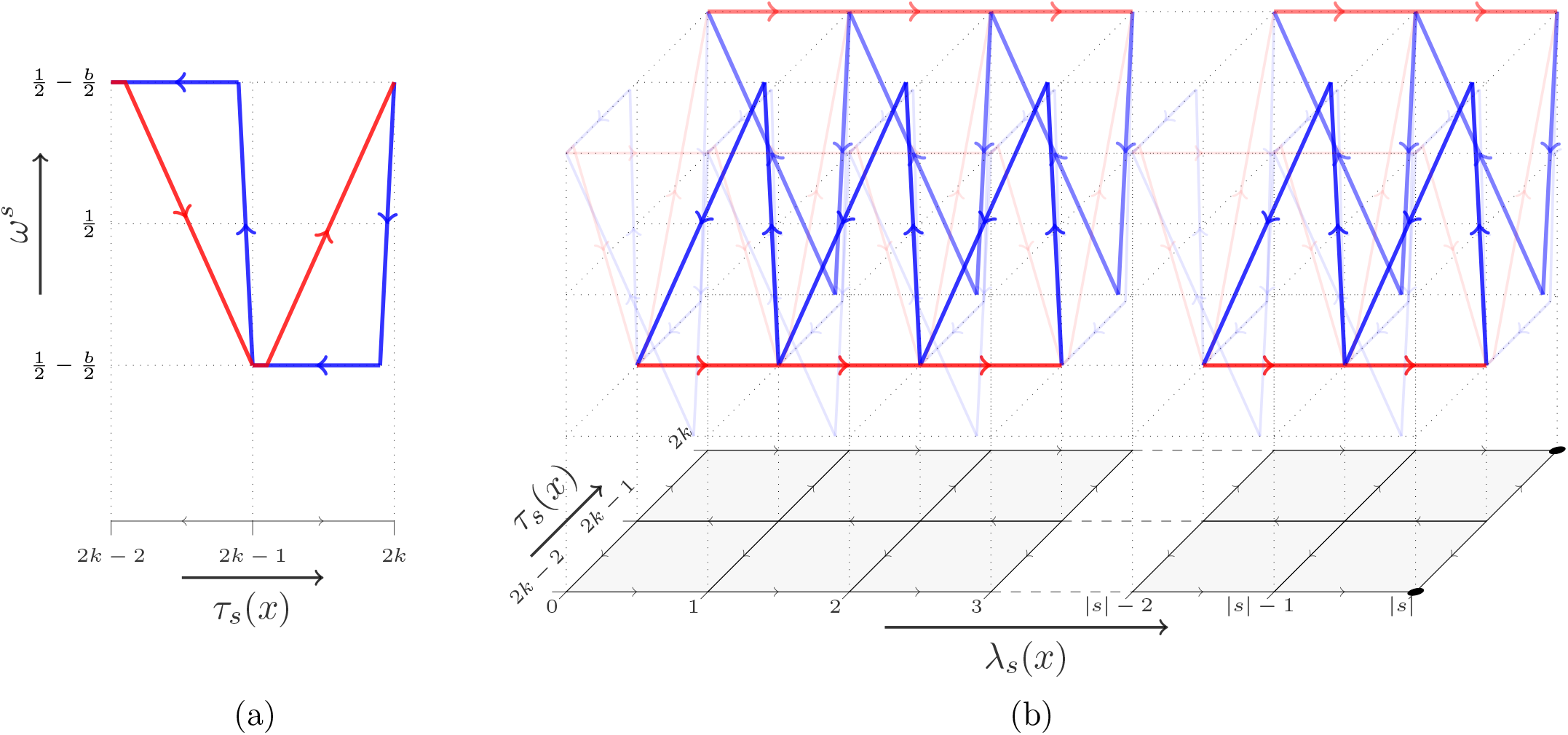
Game landscape for convergence into 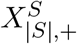 by point mutations. In each figure, genotype is shown as the base of the figure and fitness as the height above the base (i.e., *z*-axis). The base is one dimensional grid by size of tail *τ*_*s*_(*x*) in (a) and two dimensional grid by size of head *λ*_*s*_(*x*) and size of tail *τ*_*s*_(*x*) in (b). Given two adjacent discrete grid points *x*^−^ and *x*^+^ in the base, a point that is a fraction *p* between them is interpreted as a mixed population where proportion *p* has the ‘lower’ genotype *x*^−^ and proportion 1−*p* has the ‘higher’ genotype *x*^+^. The blue line shows the fitness of the lower genotype (*ω*^*s*^(*x*^−^)) and the red line shows the fitness of the higher genotype (*ω*^*s*^(*x*^+^)). Thus, if the blue line is above the red line then a mutation-limited population will tend to evolve ‘down’ towards *x*^−^ and if the red line is above the blue line then it will tend to evolve ‘up’ towards *x*^+^. This is summarized by arrow directions on the base’s grid. **(a)** shows a valley at odd-tail genotypes (*τ*_*s*_(*x*) = 1 mod 2) and peaks at even-tail genotypes (*τ*_*s*_(*x*) = 0 mod 2). This pattern repeats itself across the game landscape as *τ*_*s*_(*x*) varies from 0 to *n* − |*S*| and as *λ*_*s*_(*x*) varies from 0 to |*S*|. **(b)** takes the game landscape from (a) as the yz-plane and varies *λ*_*s*_(*x*) from 0 to |*S*| along the axis. Highlighted are the different kinds of dynamics that happen for odd-tails (front xz-plane at *τ*_*s*_(*x*) = 2*k* − 1) and even-tails (back xz-plane at *τ*_*s*_(*x*) = 2*k* − 1). For odd-tail, mutation-limited dynamics tend to decrease the size of *λ*_*s*_(*x*) and for even-tail they increase the size of the head *λ*_*s*_(*x*). In other words, the odd-tailed valleys 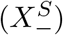 climb towards *λ*_*s*_(*x*) = 0 and the even-tailed ridges 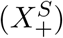 climb towards *λ*_*s*_(*x*) = |*S*|. Thus, the only local peaks under point-mutations (shown on base as black circles) have full-head and even-tail 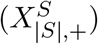.

Thus, the odd-tailed strings are not evolutionary stable under point-mutations. But more importantly, this also means that once a string becomes even-tailed, it will not go back to being odd-tailed under point-mutations. So at most one point-mutation will fix in the tail of the string.

What about mutations in the head? Consider genotypes *x* and *y* with same parity tail such that there is exactly one *i* ∈ *S* such that 0 = *x*_*i*_ *≠ y*_*i*_ = 1 (without loss of generality, by relabeling). Since *x* and *y* have the same parity tail, we have that *G*_*xx*_ = *G*_*yy*_ but *G*_*xy*_ *≠G*_*yx*_ Here there are two cases to consider: the string is still odd-tailed or it has become even-tailed.

1. For an odd-tailed string with *λτ* (*x*) = (*λ*, 2*k* − 1), we have *G*_*xy*_ = 1 > 0 = *G*_*yy*_ = *G*_*yx*_ and the game matrix:

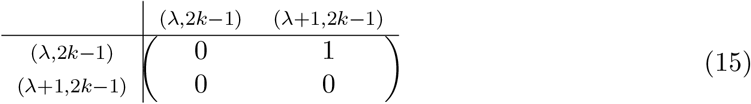

so *x* can invade *y* and thus decrease the number of 1s in the head.
2. But if *x* and *y* are even-tailed (i.e., *λτ* (*x*) = (*λ*, 2*k*)) then *G*_*xy*_ = 0 but *G*_*yx*_ = 1 giving us the game matrix:

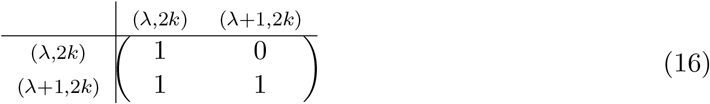

so *y* can invade *x* since (this also means, unsurprisingly, that *x* cannot invade *y*). Thus, for even-tailed strings, point mutations in the head will only fix if they increase the number of 1s in the head.

This slice of the game landscape with odd-tail valley leading up to an empty head and an even-tail ridge leading up to a full head can be seen in Figure 4b.

### Evolutionary stable set under point-mutations

From this, we can conclude that the only strings that are evolutionary stable under point-mutations have a full-head and even-tail. I will call this set 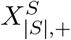. In the space of point-mutations, this evolutionary stable set is a collection of isolated local fitness peaks. More importantly, we will converge to a point in this evolutionary stable set quickly. In the worst case, it will take 2|*S*| + 1 many fixations if the population starts with a full-head, odd-tail string and then loses every 1 in the head (|*S*| fixations), flips to even-tailed, and then regains every 1 in the head. On average, it will take fewer than 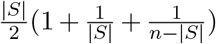 fixations.□

Although points in the full-head-even-tail set 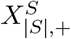 are evolutionary stable and isolated under point-mutations (i.e. strict local peaks), they are not isolated under double-mutations. Hence, after the population fixes at a point-mutation peak in 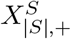, we need to focus on the dynamics of double-mutations. In terms of timescales, in a mutation limited population with mutation rate *µ*, this means switching from a timescale of 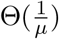 to 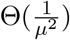.

#### Proposition 4.

*Given any parity environment f*_*n,s*_ *and any starting genotype in* 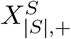, *SSWM dynamics in populations of size M* ∈ Ω(*n*^3^) *in the extended Algorithmic Darwinism framework will reach the global fitness peak at s after an expected O*(*n*) *many fixations of double-mutations and point-mutations*.

*Proof*. Let us analyze how double-mutations can move a population at genotype *x* in 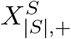. There are three kinds of bits of a string in 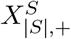: 1s in the head, 1s in the tail, and 0s in the tail. Let 2*k* be the even number of 1s in the tail. Call the double mutant genotype *y*.

### Double mutation in head

Suppose that both mutations are in the head. This happens with probability 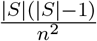. Then *G*_*yx*_ = *G*_*xx*_ = *G*_*yy*_ = *G*_*xy*_ = 1 or organized in a matrix:

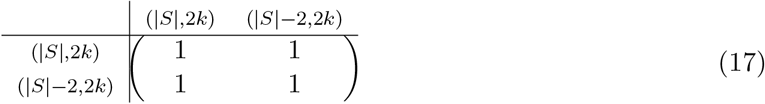

and thus *y* can only fix by random drift (i.e. with probability 1*/M*). But after it fixes, we have an even-tailed population *y* outside of 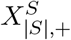 and so it will be quickly returned to 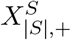 by point-mutations without changing the tail. Thus, this double-mutation does not change our position in 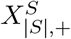. This slice of the game landscape can be seen in Figure 5a.

**Figure 5:**
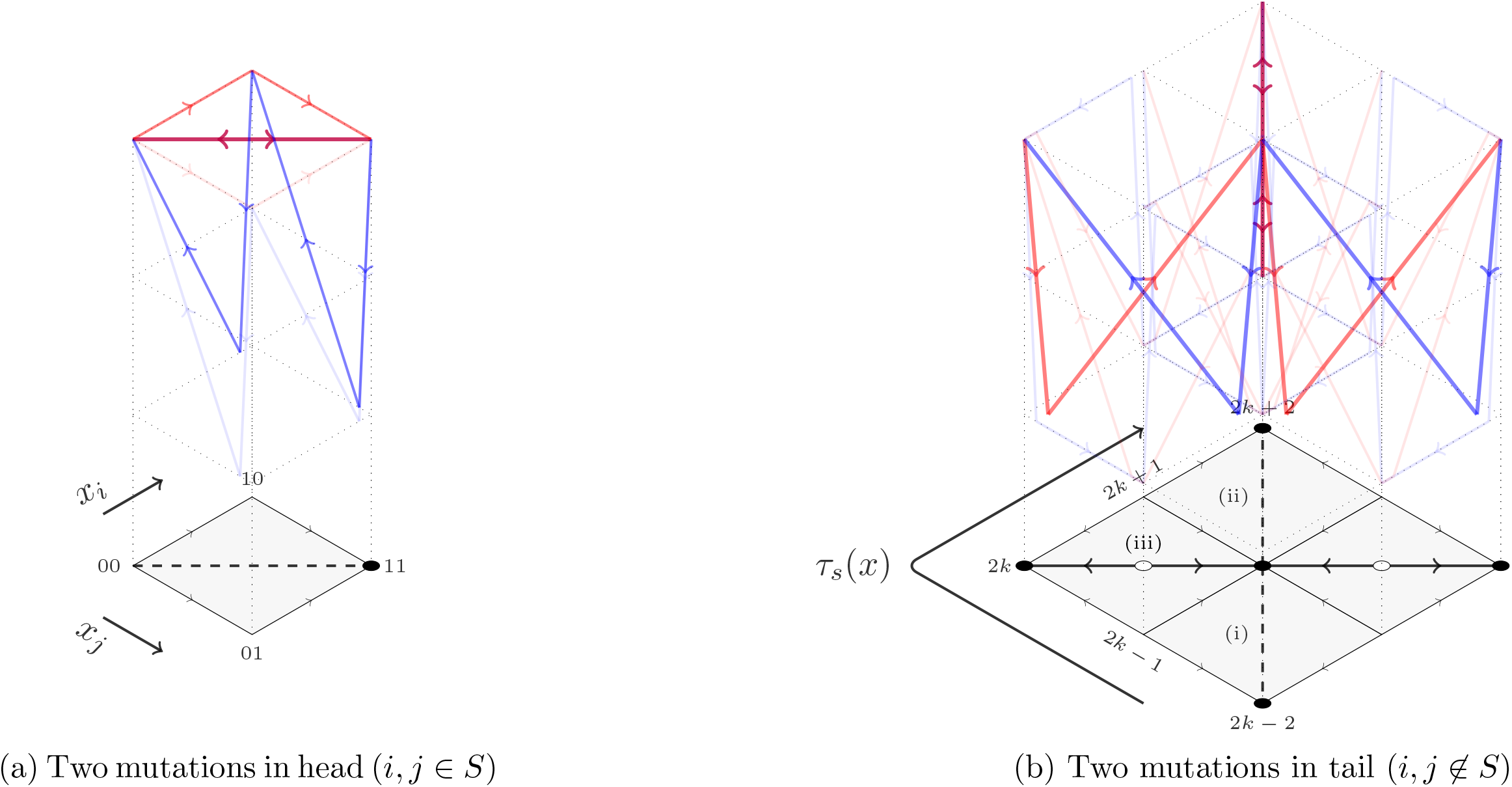
Game landscape for double-mutations in head (a) or tail (b) of full-head even-tail genotype. Base is genotypes and height is fitness as described in the caption of Figure 4. In addition to point-mutations, each figure also shows double-mutations. **(a)** shows double-mutation in the head creating an invader of equal fitness (shown as purple in game-landscape graph) that can only fix by neutral drift (shown as a dotted line in base) that is then undone by point-mutations. (b) shows three possible kinds of double-mutations in the tail: (i) both mutations flipping 1s to 0s decreasing *τ*_*s*_ by 2; (ii) both mutations flipping 0s to 1s increasing *τ*_*s*_ by 2; or (iii) one flipping a 0 to 1 and the other 1 to 0 thus maintaining *τ*_*s*_. Mutations (i) and (ii) create an invader of equal fitness that can only fix by neutral drift. Mutation (iii) creates an invader that has stag-hunt-like dynamics with resident wildtype, with invader having lower fitness at low frequency and higher fitness at high frequency creating an unstable cross in fitness. This kind of double mutant cannot invade and this dynamic is shown in base as an empty circle with arrows pointing out.

### Double mutation in tail

Consider that both mutations are in the tail. This happens with probability 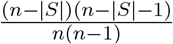. Since this preserves tail parity: *G*_*yy*_ = 1. There are three cases that produce two possible dynamics:

1. If one mutation flips 0 to 1 and the other flips 1 to 0 then *G*_*xy*_ = *G*_*yx*_ = 0 or organized in a matrix:

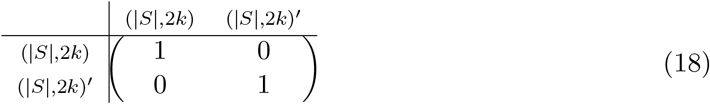

and thus *y* will be selected against and unable to fix. This happens with conditional probability 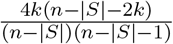
2. The remaining two cases are if both tail mutations are 0 to 1 or both are 1 to 0. These happen with conditional probabilities 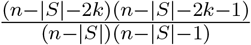 and 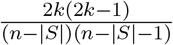, respectively. In these two cases, *G _yx_* = *G _xx_* = *G* _*yy*_ = *G*_*xy*_ = 1 or organized as a matrix:

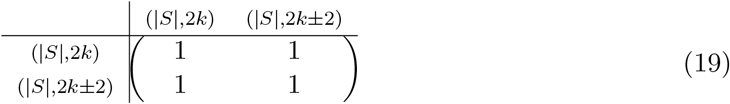

and so *y* can fix only by random drift (with probability of 1*/M*). But unlike the double-head case, the resulting *y* is still in 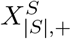 and so the fixation will not be undone by point-mutations.

This slice of the game landscape can be seen in Figure 5b.

### One mutation in head, one in tail

Finally, suppose a double mutant arises with one flip in the head and one in the tail. This happens with probability 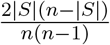 and gives us *G*_*xx*_ = 1 > 0 = *G*_*yy*_. There are two cases based on where the tail mutation hits:

1. If the tail mutation flipped a 0 to a 1 then *G*_*yx*_ = 0 with the game matrix give by:

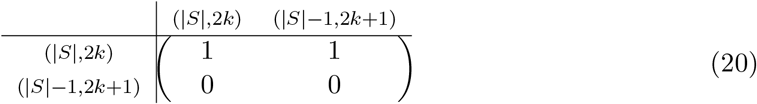

so *y* with *λτ* (*y*) = (|*S*| − 1, 2*k* + 1) is selected against and cannot fix. This happens with conditional probability 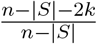.
2. The most interesting case is if the tail mutation flipped a 1 to a 0 then *G*_*yx*_ = 1 and *G*_*xy*_ = 0, giving a game matrix:

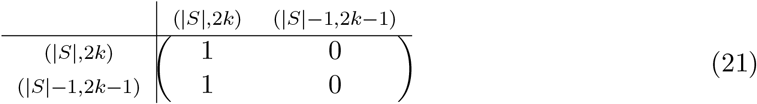 This happens with conditional probability 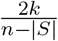. In the infinite population limit or with self-interactions, *y* can only fix by random drift. But without self-interactions in finite populations, *u*_*y*_ −*u*_*x*_ = 1*/M* so for very strong-selection (*β* > *M*), *y* will fix with probability lower bounded by a constant *ϕ* ^*^. After fixing, however, *y* is outside 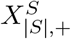 and will be quickly returned to 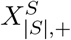 by point-mutations. Upon returning to *x*^′^ in 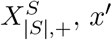 will have either one pair fewer or the same number of 1s in the tail than/as *x*. The former will happen with probability 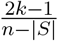 and the latter with probability 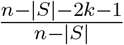.

This slice of the game landscape can be seen in Figure 6.

**Figure 6:**
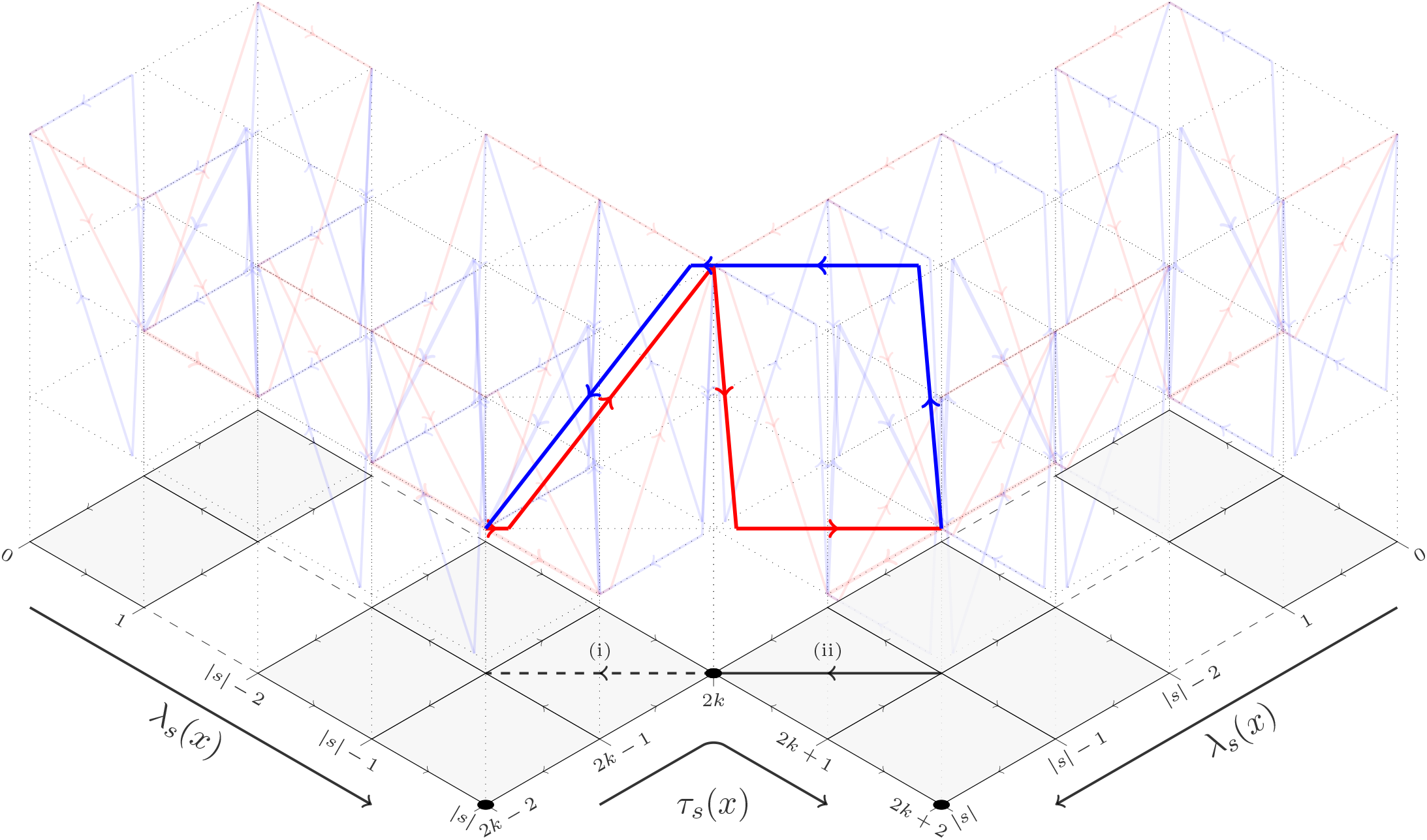
Game landscape for double-mutation with one mutation in head (*i* ∈ *S*) and one mutation in tail (*j* ∉ *S*) of a full-head even-tail genotype. Base is genotypes and height is fitness as described in the caption of Figure 4. In addition to point-mutations, there are two double mutations: (i) decreasing both head and tail, or (ii) decreasing head and increasing tail. Each wing of the figure is a copy of the game landscape from Figure 4b. Mutation (i) creates an invader of nearly-neutral fitness, with a slight fitness of advantage of 1*/M* due to the effects of a finite population of *M* individuals. In finite populations with very strong selection (*β* > *M*), this mutant fixes with probability higher than a fixed constant *ϕ* ^*^. This is represented in the base as a directed dotted line (to distinguish it from the directed solid lines that do not rely on finite populations for fitness difference). Mutation (ii) creates an invader of strictly lower fitness that cannot fix. Since both mutations (i) and (ii) lead to a point with odd-tail, the point-mutation dynamics then return the population to an even-tail of 2*k* or 2*k* − 2 (for mutation (i)), and 2*k* or 2*k* + 2 (for mutation (ii)) as described in Figure 4b.

### Biased random walk on 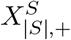

Putting this all together, we get that the double mutant regime is equivalent to a biased random walk on the integer line [0, *m*] with 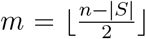 where being at a point *k* ∈ [0, *m*] represents having a genotype with a full-head and an even-tail of length 2*k*. Importantly, this walk has only a single absorbing state at *k* = 0 (i.e. when *x* = *s*). The transition probabilities for the random walk’s Markov chain are:

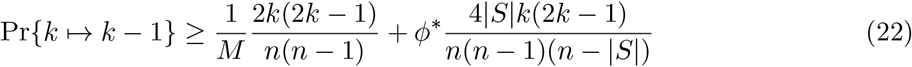

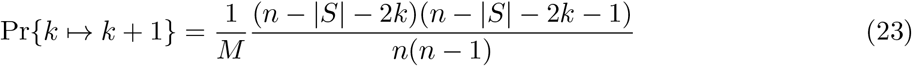

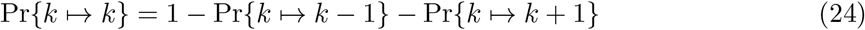

If 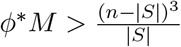 then Pr{*k* → *k* − 1} > Pr{*k* → *k* + 1} for all 1 ≤ *k* ≤ *m* − 1. So in the case that *M* = Θ(*n*^3^), the random walk will drift left, converging to *k* = 0 in number of double-mutations that is linear in the initial *k* and thus also linear in *n* − |*S*|.□

Now we can put together Propositions 3 and 4 to get the total adaptation time measured in either fixations or Moran process birth-death events:

#### Proposition 5.

*SSWM dynamics in the extended Algorithmic Darwinism framework will find the unique fitness peak s of any parity environment f*_*n,s*_ *after* 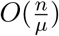 *many mutations and O*(*n*^7^) *birth-death events*.

*Proof*. Let us first look at the number of mutations and then translate this to the number of birth-death events.

### Number of mutations

Putting together Propositions 3 and 4 this results in an overall *O*(|*n*|) number of fixations with evolution proceeding fast then slow. The fast part proceeds via a linear number of point-mutations with each on a time-scale of 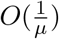. The slow part proceeds by a linear number of double-mutations, but since each double-mutation arises on the time-scale of 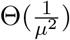, we will expect to see a total of 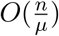 point-mutations. Of all those point-mutations, most will not fix – the exception is the *O*(*n*) point-mutations that follow the one-mutation-in-head-one-in-tail double-mutation. Thus, I expect on the order of 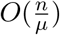 total mutations.

### Number of birth-death events

To finish the time analysis, we can go past mutations to the number of birth-death events that make up each Moran process. For all the mutations that fail to fix, they will fail with an expected *O*(1) number of birth-death events. Of the mutations that do fix: the ones that are selected for will fix in an expected *O*(*M* ln *M*) number of birth-death events and the ones that are neutral will fix in an expected *O*(*M* ^2^) number of birth-deaths [40]. Since invasion by neutral drift happens only for the double mutants, this means that the overall evolutionary time can be bounded by 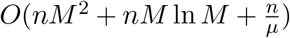. But we need to avoid a new mutation arriving while an existing one is fixing, so I need to set 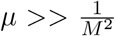 and from Proposition 4 we need *M* = Θ(*n*^3^). Thus, the overall evolutionary time measured in the number of birth-death can be bounded by *O*(*n*^7^).

Of course, *O*(*n*^7^) is a long time and I expect that future work will be able to improve this bound. But, it is already a polynomial and not an exponential bound. This polynomial bound is sufficient to show that an evolving population can adapt to the Parity environment under extended algorithm Darwinism and thus prove Theorem 1.

## 6 Discussion

Theoretical computer science and evolutionary biology provide a rich interface for the development of new theory. Providing a formal model of computation that characterizes which algorithms correspond to evolution is essential to this new interface. Valiant [1] started by defining strict algorithmic Darwinism as a subset of SQ-learning, but this did not account for the rich feedback possible between ecology and evolution [11, 12] through effects like frequency-dependent selection. By extending algorithmic Darwinism to include these ecological interactions, I showed that evolution is computationally more powerful than we previously suspected. Thus, we need to continue to refine our model of evolution to better integrate theoretical computer science into biology.

The hardness of Parity in strict algorithmic Darwinism comes from the fact that, when an organism encounters a specific challenge, it has no way of communicating the content of that specific challenge to its progeny. The details of specific challenges are lost in the statistic of fitness. In the extended algorithmic Darwinism, I allow a way around this. Organisms convey some information about the challenges because the resident organisms provide specific challenges to the invader. This bias in the distribution makes extended algorithmic Darwinism a subset of what is learnable with extended Statistical Queries (or, equivalently PAC + membership queries) [14]. In fact, given that noisy-Parity is conjectured to not be PAC-learnable [26, 27], it is reasonable to believe that the full frequency-dependent distribution of challenges of the extended Algorithmic Darwinism cannot be simulated with just example queries. The frequency-dependent distribution of challenges can, however, be simulated with extended statistical queries [14] or a combination of example and membership queries (MQ). This raises the open question of where exactly to locate extended algorithmic Darwinism between PAC and PAC + MQ.

Of course, frequency-dependence is not the only way that extra information could be transferred between generations. For other mechanisms, organisms could encode some of the examples into their DNA (epigenetics), or some organisms might programmatically duplicate certain examples in the world to adjust their relative frequency for their offspring (niche construction and niche inheritance). These are other ways that future work could extend algorithmic Darwinism to give us a clearer picture of how these various mechanisms – that are often treated as extra flourishes by the modern evolutionary synthesis – can contribute to the fundamental computational power of evolution.

### Prototrophs vs heterotrophs and the origin of life

In the context of the Oparin-Haldane hypothesis for the heterotrophic origin of life, the organisms in both strict and extended algorithmic Darwinism would, technically, be classified as heterotrophs. This is because these organisms do not generate their own energy but have to harvest it from correctly consuming macromolecules in the environment. But there is also a fundamental difference between these two kinds of organisms. In strict algorithmic Darwinism, all the macromolecules are generated by the environment in some process that is independent of the organisms in that environment. This sort of metabolism is exceptionally rare on Earth, maybe occurring only in deep-sea hydro-thermal vents. As such, I think that the organisms of strict algorithmic Darwinism should be called *prototrophs* to distinguish them from typical heterotrophs.

The *heterotrophs* that do occur frequently in nature are much more similar to the organisms of the extended algorithmic Darwinism, in that they harvest energy from correctly eating macro-molecules that are generated by a process that depends on the distribution of organisms in the environment. But it is exactly this shift from an abiotic to biotic environment that I have shown to transform the mode of evolution to make it exponentially more powerful. Thus, it is tempting to hypothesize that part of the reason that we have such an abundance of true heterotrophs from a presumably prototropic origin is due to the computational resource that heterotrophy gave populations in adapting to Earth’s early environment.

More generally, we can also consider the evolution or proliferation of predation as an important driver of frequency-dependent selection. As noted by Bengtson [41]: “predation played a crucial role in some of the major transitions in evolution” including the origin of eukaryotes, multicellularity, animal motility, and the vast proliferation of kinds during the Cambrian explosion. Given the computational resources that frequency-dependent selection unlocks for evolution, maybe the origin of frequency-dependent selection or the origin of predation should be considered as its own major transition in evolution.

Many consider the ability to metabolize, replicate and evolve as the central characteristics of living organisms. We should augment this list: the central characteristics of living organisms are the ability to metabolize, replicate, evolve and interconnect in a shared ecology.

### Ecology is more transformative than sex

It is also important to highlight just how transformative ecology is to the mode of evolution. For example, by comparing it to the difference in mode between asexual vs sexual reproduction. Sexual reproduction, recombination, horizontal gene-transfer, and fusion are another set of processes that are believed to have the potential to greatly speed-up evolution [42–44] and even matter in the origins of life [45] and the somatic evolution of cancer [46]. Strict algorithmic Darwinism has previously been expanded to study the transformation from asex to sex by considering recombination [47] and horizontal-gene transfer [48, 49]. The authors showed that an exponential speed up is possible in the number of generations required for adapting-to a family of easy environments. But unlike my results for ecology, they also proved that the set of families that are adaptable-to is not changed and is still the set of CSQ-learnable environments.

In contrast, in this paper, I presented an exponential speed-up for the Parity environment that is hard for strict algorithmic Darwinism. This moves the ecologically extended algorithmic Darwinism beyond what is CSQ-learnable. The sexual mode of evolution allowed a speed-ups from polynomial to polylog number of generations. But, unlike the ecological mode of evolution, the sexual mode of evolution did not allow for a speed-up from exponential to polynomial number of generations. In other words, both sex and ecology can dramatically speed up evolution, but it is only ecology that can transform what kind of environments are adaptable-to. This transformation in computational power might encourage us to re-prioritize the central tenets of evolution from ‘selection, mutation, and recombination’ to ‘selection, mutation, and interaction’.

Knowing about this increase in computational power is important not only for a general understanding of evolution, but also so that we do not underestimate the power of eco-evolutionary dynamics in settings where evolution is our adversary. For example, in cancer, the patient and physician are in a battle against somatic evolution [50, 51]. Since we tend to view the change in gene-frequencies as the most important aspect, we tend to treat cancer as a disease of individual aberrant cells. But given that frequency-dependent selection can so drastically transform the mode of evolution, we need to also focus on the aberrant ecology of cells [33, 34, 46, 50–52]. That means, that if we want to design drugs that do not allow for the evolution of resistance, we cannot have these drugs produce fitness landscapes that are hard for just strict algorithmic Darwinism but instead we need to design drugs that produce game landscapes that are hard for the ecologically extended algorithmic Darwinism. My hope is that a formal algorithmic theory of eco-evolutionary dynamics like my ecologically extended algorithmic Darwinism can help in this effort.

## Acknowledgements

I am grateful to Tyler Cassidy, Emily Dolson, Patrick Ellsworth, Nathan Farrokhian, Sergio Graziosi, Peter Jeavons, Marco Lin, Jeff Maltas, Rob Noble, Georgios Piliouras, Thatchaphol Saranurak, and Julian Xue for feedback on an earlier draft of this article; and to Rahul Santhanam for helpful discussion. The Darwinian engine in Figure 1 was inspired by presentations given by Amitabh Joshi and Joachim Krug. This research was supported by a fellowship from the James S. McDonnell Foundation.

## Notes

### Competing Interest Statement

The authors have declared no competing interest.

### Summary of Updates

Main results restructured as theorems and proofs. Several figures added for clarity. In particular, I added a number of visual representations of the Parity game landscape.

